# Trait plasticity and covariance along a continuous soil moisture gradient

**DOI:** 10.1101/2020.02.17.952853

**Authors:** J. Grey Monroe, Haoran Cai, David L. Des Marais

**Affiliations:** Department of Plant Sciences, Unive*f*rsity of California at Davis, Davis, USA; Department of Civil and Environmental Engineering, Massachusetts Institute of Technology, Cambridge, MA, USA; The Arnold Arboretum of Harvard University. Boston, MA, USA; Max Planck Institute for Developmental Biology, Tubingen, DE

**Keywords:** drought, function-valued traits, Brachypodium, evolutionary constraint, phenotypic plasticity, non-linearity

## Abstract

Water availability is perhaps the greatest environmental determinant of plant yield and fitness. However, our understanding of plant-water relations is limited because it is primarily informed by experiments considering soil moisture variability at two discrete levels – wet and dry – rather than as a continuously varying environmental gradient. Here we used experimental and statistical methods based on function-valued traits to explore responses to a continuous soil moisture gradient in physiological and morphological traits in two species and five genotypes each of the model grass *Brachypodium.* We find that most traits exhibit non-linear responses to soil moisture variability. We also observe differences in the shape of these non-linear responses between traits, species, and genotypes. Emergent phenomena arise from this variation including changes in trait correlations and evolutionary constraints as a function of soil moisture. These results point to the importance of considering non-linearity in plant-water relations to understand plastic and evolutionary responses to changing climates.

## Introduction

For plants, soil water availability is an important environmental factor in ecology and agriculture, acting as a major determinant of fitness and yield (Juenger 2013; Greenham *et al*. 2017). As such, considerable effort has been placed on studying plant responses to drought, often defined conceptually and experimentally as an environmental condition of abnormally elevated aridity resulting in decreased plant performance (Passioura 1996). Most of the research on drought responses and tolerance strategies, including in *Brachypodium,* the focal system of this work, has involved comparisons between discrete soil water levels – control and water-limited (Des Marais *et al*. 2012; Edwards *et al*. 2012; El-Soda *et al*. 2014; Vasseur *et al*. 2014; Greenham *et al*. 2017). Yet soil moisture as an environmental factor is complex and multidimensional, with fluctuations varying continuously in timing, duration, and degree. Here we investigate trait responses to one important dimension of soil moisture variability – degree – with experimental and statistical approaches treating soil moisture content as a continuous variable rather than a set of fixed levels.

Quantitative genetic variation in drought resistance traits have been observed in natural populations and laboratory model systems. In particular, natural populations of Brassicaceae species (including *Arabidopsis thaliana)* and *Brachypodium* harbor variation both constitutive and plastic traits mediating plant-water relations, including water use efficiency (Des Marais *et al*. 2012, 2017; Edwards *et al*. 2012; Greenham *et al*. 2017), leaf chemistry (Kesari *et al*. 2012; Des Marais *et al*. 2017), leaf anatomy (Skirycz *et al*. 2011; Verelst *et al*. 2013; Dittberner *et al*. 2018), root-shoot biomass partitioning (Des Marais *et al*. 2012, 2017), and many others (Verslues and Juenger 2011; Edwards *et al*. 2012; Juenger 2013; Luo *et al*. 2016; Yarkhunova *et al*. 2016; Lenk *et al*. 2019). Among the many plant traits that can and have been measured, few have been studied more extensively than specific leaf area (SLA – often reported as its inverse, leaf mass per area or LMA). SLA provides a description of leaf architecture that is central to the leaf economics spectrum, a theory which seeks to explain variation in leaf physiological strategies, from more conservative (low SLA) to more productive (high SLA) (Wright *et al*. 2004). In the context of drought stress, low SLA might be adaptive as lower leaf surface area is expected to reduce water loss through transpiration. In several cases, reductions in SLA have been reported under drought conditions (Casper *et al*. 2001). However, for all of these traits, the shape of plastic responses to variation in water availability remain largely unmeasured.

Less is known about how trait variation and relationships between traits change across continuous environmental gradients (but see Robinson *et al*. 2009). Genetic variation for traits can be higher, for example, in less frequently encountered environmental conditions (Gibson and Dworkin 2004; Schlichting 2008; Paaby and Rockman 2014). And describing this structure is important as trait variances and covariances influence evolutionary constraints. For example, if total genetic variance in trait space changes depending on the environment, then the capacity to respond to selection will vary accordingly, with reduced responses to selection under conditions where genetic variance is lower and vice versa. Similarly, if trait covariances depend on environment (Wood and Brodie III 2015), conditions which increase trait covariances may limit evolutionary potential across a range of environments by reducing the effective axes of variation (Levins 1968; Lande and Arnold 1983; Via and Lande 1985; Kingsolver and Gomulkiewicz 2003; Gomulkiewicz *et al*. 2018). These phenomena are made more complex by the possibility that the relationship between trait variances and covariances with the environment may be non-linear. Investigating the genetic architecture of multiple traits is therefore useful for understanding evolution in rapidly changing environments. Fortunately, this need has been met with the development of statistical methods to reduce the dimensionality of genetic variance covariance matrices and produce meaningful summaries describing evolutionary constraints of multiple phenotypes simultaneously (Houle 1992; Blows 2007; Kirkpatrick 2009; Kingsolver *et al*. 2015).

To study the shape of phenotypic responses to the environment, modeling traits as mathematical functions of continuous variables, or function-valued-traits, has proven powerful (Kirkpatrick and Heckman 1989; Kingsolver *et al*. 2001; Griswold *et al*. 2008; Stinchcombe *et al*. 2012; Goolsby 2015; Gomulkiewicz *et al*. 2018). Recent decades have seen this approach used across many organisms (Gibert *et al*. 1998; Pettay *et al*. 2008; Rocha and Klaczko 2012; Mason *et al*. 2020), traits (Robinson *et al*. 2009; McGuigan *et al*. 2010; Stinchcombe *et al*. 2010), and components of the environment (Brommer *et al*. 2008; McGuigan 2009; Pearse *et al*. 2019). For plant scientists, experimental and analytical approaches leveraging the concept of function-valued-traits provide a framework for gaining a deeper understanding of plant acclimation and evolutionary adaptation to the environment.

In this study we combined these approaches into an exploratory investigation of trait plasticity, covariance and evolutionary constraints across soil moisture gradients. We model trait responses to a continuous gradient of soil moisture for multiple genotypes of two species of the model grass genus *Brachypodium:* annual *B. distachyon* and perennial *B. sylvaticum.* Because the shapes of trait responses cannot necessarily be known *a priori*, we use model selection among linear and non-linear environmental predictors to estimate the response function for each trait. We then estimate genotype means at different levels of soil moisture and compute the variance-covariance parameters for all traits. Finally, we ask whether patterns of variance and covariance in drying-responsive traits in *Brachypodium* species may lead to variation in evolutionary constraint as a function of soil moisture.

## Materials and Methods

### Genotypes and species

*Brachypodium* is a model genus for the genetics and genomics of C3 grasses (Brkljacic *et al*. 2011). In this work we studied natural variation between and among two species of *Brachypodium:* the annual *B. distachyon* and the perennial *B. sylvaticum.* Both species are endemic to Eurasia, with *B. distachyon* more prevalent in seasonally dry habitats in Southern Europe, North Africa, and the Middle East, and *B. sylvaticum* more widely distributed throughout Eurasia (Catalan *et al*. 2016) (Figure S1). The different evolutionary histories, ecological environments, and life habits of these two species may have impacted patterns of genetic variation and evolvalbility in relation to soil moisture.

To investigate genetic variation, five genotypes of each were studied to characterize patterns of variation in plant traits across an environmental gradient: *Brachypodium distachyon* inbred lines ABR2, Adi-10, Bd21, Bd3-1, and Koz-1 and *Brachypodium sylvaticum* inbred lines Ain-1, Ast-1, Kry-1, Osl-1, Vel-1. For each species, these genotypes represent a range of geographical origins and phenotypic diversity (Personal observations) (Steinwand *et al*. 2013; Des Marais *et al*. 2017). For example, Kry-1 is native to Krym, Ukraine while Ain-1 is native to Ain Draham, Tunisia (Steinwand *et al*. 2013). Both species are self-compatible and each of the lines used here have been maintained as inbred lines for greater than six generations (Vogel *et al*. 2009; Steinwand *et al*. 2013); as such, experimental replicates may be considered nearly isogenic.

### Plant growth and dry down experiment

Plant growth and experimental soil dry down were performed in the greenhouses of the Arnold Arboretum of Harvard University. To synchronize germination across genotypes within each species, seeds were placed on damp filter paper in the dark at 4°C for 14 days prior to planting. To synchronize the developmental stage at the timing of the drought treatments between the two species, *B. sylvaticum* seeds were planted thirteen days before *B. distachyon* (Oct 7 and 20, 2015, respectively). For each genotype, 1200 seeds were planted two to a pot and were subsequently thinned to one plant, for a total of 600 experimental plants in a randomized block design. All plants germinated within four days of sowing. Individual seeds of plants were sown in Greens Grade Profile porous ceramic rooting media (Profile Products, Buffalo Grove, IL, USA) in Deepot D40H Conetainers (650mL; Stuewe & Sons. Tangent, OR, USA) and grown at 25°C/20°C days/nights. Ambient sunlight was supplemented to 1000 umol/m2/s for 12hr/day.

Dry down treatments began 29 and 42 days after sowing (DAS) for *B. distachyon* and *B. sylvaticum,* respectively. Because the harvesting was divided over five consecutive days (see section below), plants were split into five equal harvest cohorts, with each cohort containing equal numbers of each watering treatment to avoid confounding harvest day with soil moisture content. Thus, though each consecutive cohort differed in age by a single day, each experienced the dry down treatment for the same amount of time. Nevertheless, we expected the difference in age between harvest cohorts to potentially impact trait expression and we therefore included harvest day (cohort) as a covariate in subsequent models. To generate a continuous gradient of final soil moisture by the end of the dry down period, plants were split into five watering treatments, receiving approximately 0,4, 8, 12, 16, or 20 ml of water per day for 14 days (Figure S2). Prior to initiating the experiment pots were weighed with dry soil (*massdry*) and field capacity *(mass_max_*). During the course of the dry down, soil moisture content was calculated during the morning of day *d* for each pot as *mass_d_*/(*mass_max_ – mass_dry_*). The final soil moisture reflects the combined effects of water input and output through evaporation from the soil and evapotranspiration from the plants.

### Plant harvesting and phenotyping

To characterize phenotypic responses to our experimental soil moisture gradient, we measured a suite of developmental and physiological traits. Plants were harvested in five cohorts over five days at the end of the dry down period. Each day, half of the sampled plants were harvested for above and below ground biomass, total above ground green area, *δC*_13_, N content, C content. The other half were harvested and assessed for specific leaf area (SLA) and relative water content (RWC). Above ground leaf area was estimated by laying freshly harvested plants flat between plates of clear acrylic and imaging with a Nikon 5300 digital camera at a fixed distance with a 35mm Nikkor lens. Total green pixels were counted for each image with Easy Leaf Area (Easlon and Bloom 2014) (https://github.com/heaslon/Easy-Leaf-Area) with settings shown in Figure S3. Above ground biomass was measured after drying leaf material overnight at 60°C and then for several weeks at room temperature. Below ground biomass was measured after washing the soil matrix from roots and drying them overnight at 60°C and then for several weeks at room temperature. Above and below ground biomass was measured after leaves and roots were dried. Leaf tissues for *δC*_13_, *δN*_15_, nitrogen (hereafter “N”) content, and carbon (hereafter “C”) content were ground to a fine powder and processed by the UC Davis Stable Isotope Facility. SLA was calculated by scanning the two youngest fully emerged leaves with a 1 *cm*^2^ red square. Leaf area in *mm*^2^ was calculated from these same images using Easy Leaf Area. SLA was calculated as *leaf area/biomass_dry_*. These leaves were also used to calculate RWC. Prior to drying, fresh leaves were weighed (*biomass_fresh_*) and then submerged under water in 15mL falcon tubes for several hours. They were then weighed *(biomass_fresh_*), oven-dried overnight, and weighed again (*biomass_dry_*). RWC was calculated as (*biomassf_resh_ – biomass_dry_*)/(*biomass_turgid_ – biomass_dry_*).

## Analyses

We used R for all statistical analyses. Code and data to generate this manuscript can be found at https://github.com/greymonroe/brachypodium_fvt and Zenodo (10.5281/zenodo.4446263).

### Function-valued traits

For the purposes of modeling phenotypic responses to variation in soil moisture content, we considered soil moisture content as the final soil moisture on day 14 of the dry down period for each plant, referred to in figures as *S oil moisture(%_final_*). A major challenge in studying function-valued traits is model selection. That is, identifying the functions that best describe the curvature (or lack thereof) in the shape of phenotypic responses to environmental gradients. Quadratic and natural splines have been suggested as potential functions to model non-linearities (Meyer 2005), but assuming the appropriate function is problematic. Akaike information criterion (AIC) selection based on contrasting multiple complex models offers an effective means to balance predictability with over-fitting (Griswold *et al*. 2008; Bolstad *et al*. 2010; Gomulkiewicz *et al*. 2018). Because we sampled only 5 genotypes per species (Van Eeuwijk 1995) we treat genotype as a fixed effect in our model, which differs from random regression used elsewhere where genotypes are treated as random effects (Nussey *et al*. 2007). Thus, we began with the full fixed-effects linear regression model for traits as described below.

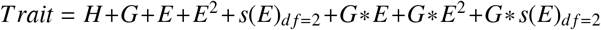

where *H* = *harvest day, G* = *genotype, E* = *soil moisture, E*^2^ = *quadratic parameter, s(E)*_*df*=2_ = *second degree natural spline parameter*

We then selected a model for each trait using stepwise AIC model selection with the *stepAIC* function from the package *MAS S* (Venables and Ripley 2002) in R with the “direction” parameter set to “both.” The two species were analyzed separately to avoid biases introduced by enforcing the same model on species with different sizes, developmental trajectories and evolutionary histories.

### Genetic correlations as a function of soil moisture content

We calculated trait correlations at different levels of soil moisture to characterize how genetic correlations between traits vary as a function of soil moisture content. Predicted genotypic means for each trait were calculated at 20 levels of soil moisture content (from 30% to 100% gravimetric water content) based on the model chosen by AIC (see above). Pearson correlation coefficients between genotype means were calculated at each level of soil moisture within each species.

### Plasticity through multidimensional trait space

We quantified total plasticity through multidimensional trait space as a function of soil moisture by first scaling each trait to a mean of 0 and standard deviation of 1. We then calculated the euclidean distance matrix between genotype means at all soil moisture levels. We then estimated total plasticity, measured as euclidean distance between trait values, between consecutive soil moisture levels for each genotype. *m* is a vector of length 20 evenly spaced soil moisture measurements from 30% to 100%:

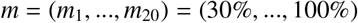

If *p_g,i_* = (*p*_*g*,1_,...,*p_g,n_*) and *p*_*g,i*+1_ = (*p*_*g,i*+1,1_,...,*p*_*g,i*+1,*n*_) are two consecutive (in terms of soil moisture) points in *n* dimensional trait space of scaled genotype means, total plasticity (Euclidean distance) for genotype *g* across *n* traits at soil moisture level *i* is calculated as

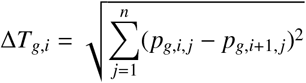

Finally, to ask if total plasticity differed between the two species, at each level of soil moisture, we then compared the mean Δ*T* between *B. sylvaticum* and *B. distachyon* by T-tests. To visualize plasticity of each genotype through multivariate trait space further, we performed a principal component analysis from the matrix of scaled genotype trait means using the *prcomp* function in R.

### Analysis of evolutionary constraints among traits

We calculated several statistics summarizing evolutionary constraint as described in Kirkpatrick (2009). First, for each species, we approximated the *G* matrix of genetic covariances by calculating variances and covariances between mean scaled genotype trait means at different levels of soil moisture. We then calculated, using the *prcomp* function in R, the eigenvalues of each mean standardized (trait values divided by mean) *G* matrix, *λ_i_*. From these we then calculated the *number of effective dimensions, n_D_*, equal to the sum of the eigenvalues divided by the largest eigenvalue:

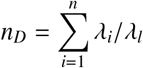

We also calculated the *maximum evolvability, e_max_*, equal to the square root of the largest eigenvalue, *λ_l_* (Houle 1992; Kirkpatrick 2009):

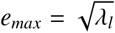

Finally, we calculated the *total genetic variance* (Kirkpatrick 2009), equal to the sum of the eigenvalues of *G*:

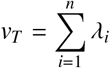

## Results

### The dry down experiment resulted in a continuous soil moisture gradient

Across the six watering treatments, combined with random variation in water capacity of pots (Figure S4), the dry down period resulted in a continuous environmental gradient of final soil moisture, albeit with a higher frequency of plants near the driest extreme of soil moisture variation (Figure 1). This gradient provides the basis for analyzing phenotypes in relation to soil moisture treated as a continuous gradient rather than limited set of discrete factors.

**Figure 1.**
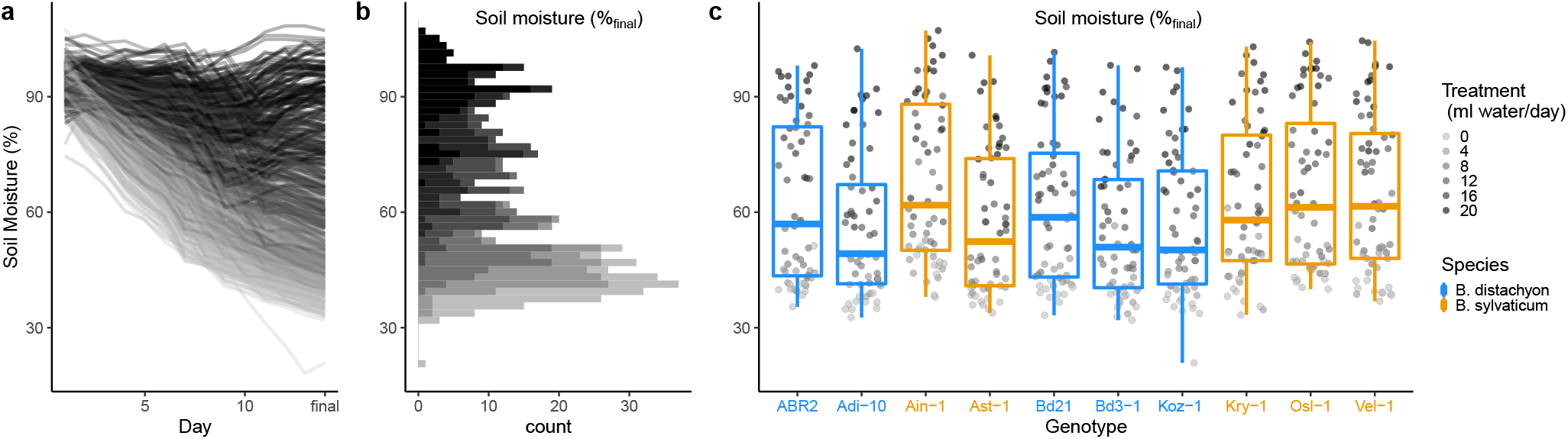
Effect of the experimental dry down on soil water content. (A) Time series of gravimetric soil moisture for all pots during the 14-day dry down period. (B) Distribution of final (day 14) soil moisture content across all pots. The data are distinguished by color according to the watering treatment. (C) Final soil moisture content by genotype.

The observed reduction in leaf relative water content (RWC) under the driest conditions in both *B. distachyon* and *B. syl-vaticum* indicates that at this extreme, plants were physiologically stressed (Figure 2A). Mean leaf RWC for plants in the 10% tail of soil water content was 85.21% which is drier than that observed in the less severe soil drying conditions of Des Marais et al. (Des Marais *et al*. 2017). Additional visual observations made during the experiment such as leaf rolling, another symptom of dehydration stress, were evident in plants at the lowest water treatment by the end of the dry down period.

**Figure 2.**
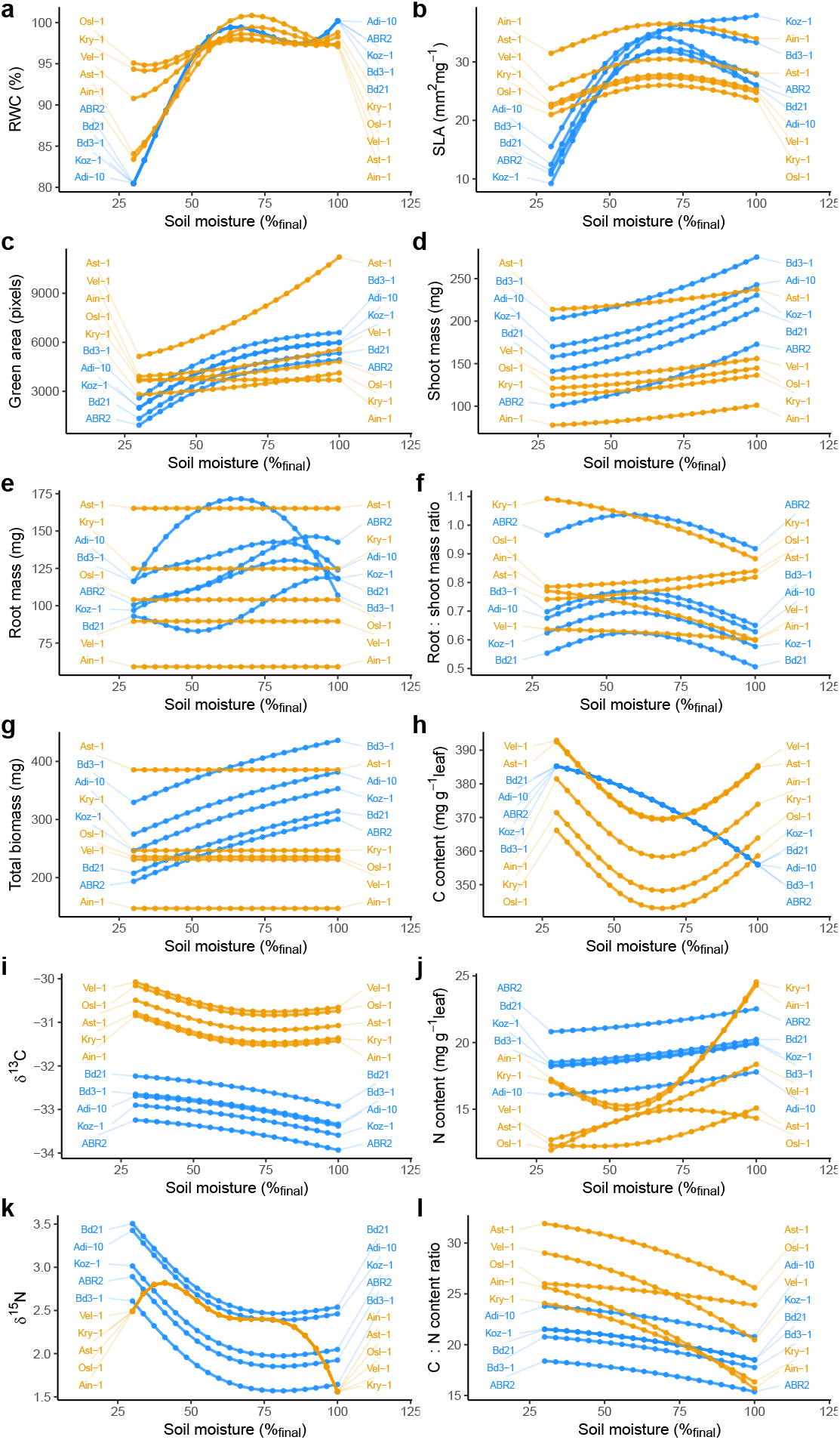
Variation in phenotypic responses to soil moisture gradient as function-valued traits. Points and the lines connecting them indicate predicted values at different levels of gravimetric soil moisture content from 30% to 100%. *B. sylvaticum* genotypes are colored orange and *B. distachyon* blue. (a) RWC (b) SLA (c) Green Area (d) Shoot Mass (e) Root Mass (f) Root:Shoot (g) Biomass (h) C content (i) d13c (j) N content (k) d15n (l) C:N ratio

### Non-linearity in trait responses to soil moisture is common

We evaluated the degree to which traits showed linear or non-linear responses to soil moisture using an AIC model selection approach from a full model which included quadratic and natural spline parameters relating soil moisture content to plant phenotypes. We observed non-linear components (quadratic, spline, or both) in the final models for all of the traits which included an environmental (water content) predictor (Table 1, Figure 2, Table S1, Table S2). In *B. distachyon,* all of the traits included at least one non-linear environmental predictor. In contrast total biomass and root mass did not include an environmental predictor in selected models in *B. sylvaticum.*

**Table 1.** Model selection. ‘x’ indicates parameters chosen in best minimum AIC model. H = harvest day, G = genotype, E = final soil moisture, E*2 = quadratic parameter, s(E) = 2nd degree natural spline parameter

|  | G | E | E<sup>2</sup> | s(E) | H | G*E | G*E<sup>2</sup> | G*s(E) |
| --- | --- | --- | --- | --- | --- | --- | --- | --- |
| RWC - B. distachyon |  |  | x | x | x |  |  |  |
| RWC - B. sylvaticum | x |  | x | x |  |  |  | x |
| SLA - B. distachyon | x |  | x | x | x |  | x |  |
| SLA - B. sylvaticum | x |  |  | x |  |  |  |  |
| Green Area - B. distachyon | x |  |  | x | x |  |  |  |
| Green Area - B. sylvaticum | x |  | x |  | x |  | x |  |
| Shoot Mass - B. distachyon | x |  | x |  | x |  |  |  |
| Shoot Mass - B. sylvaticum | x |  | x |  | x |  |  |  |
| Root Mass - B. distachyon | x |  | x | x | x |  |  | x |
| Root Mass - B. sylvaticum | x |  |  |  |  |  |  |  |
| Root:Shoot - B. distachyon | x |  |  | x | x |  |  |  |
| Root:Shoot - B. sylvaticum | x |  | x |  | x |  | x |  |
| Biomass - B. distachyon | x |  |  | x | x |  |  |  |
| Biomass - B. sylvaticum | x |  |  |  | x |  |  |  |
| C content - B. distachyon |  |  | x |  |  |  |  |  |
| C content - B. sylvaticum | x |  |  | x |  |  |  |  |
| d13c - B. distachyon | x |  | x |  | x |  |  |  |
| d13c - B. sylvaticum | x |  |  | x | x |  |  |  |
| N content - B. distachyon | x |  | x |  | x |  |  |  |
| N content - B. sylvaticum | x |  |  | x |  |  |  | x |
| d15n - B. distachyon | x |  |  | x | x |  |  |  |
| d15n - B. sylvaticum |  |  | x | x | x |  |  |  |
| C:N ratio - B. distachyon | x |  | x |  | x |  |  |  |
| C:N ratio - B. sylvaticum | x |  | x |  |  |  | x |  |

When considering plasticity across multidimensional trait space, it appears that most of the variation in *B. distachyon* is attributable to responses to low soil moisture (Figure 3, Figure S5). In contrast, *B. sylvaticum* was more responsive to extreme wet conditions than *B. distachyon.* We observed, particularly in *B. sylvaticum*, that phenotypes were similar between extreme dry and extreme wet soil moisture contents (Figure S5b). This similarity may be explained by the quadratic parameters of trait functions where the curvature of trait responses can lead to similar phenotypes at both environmental extremes.

**Figure 3.**
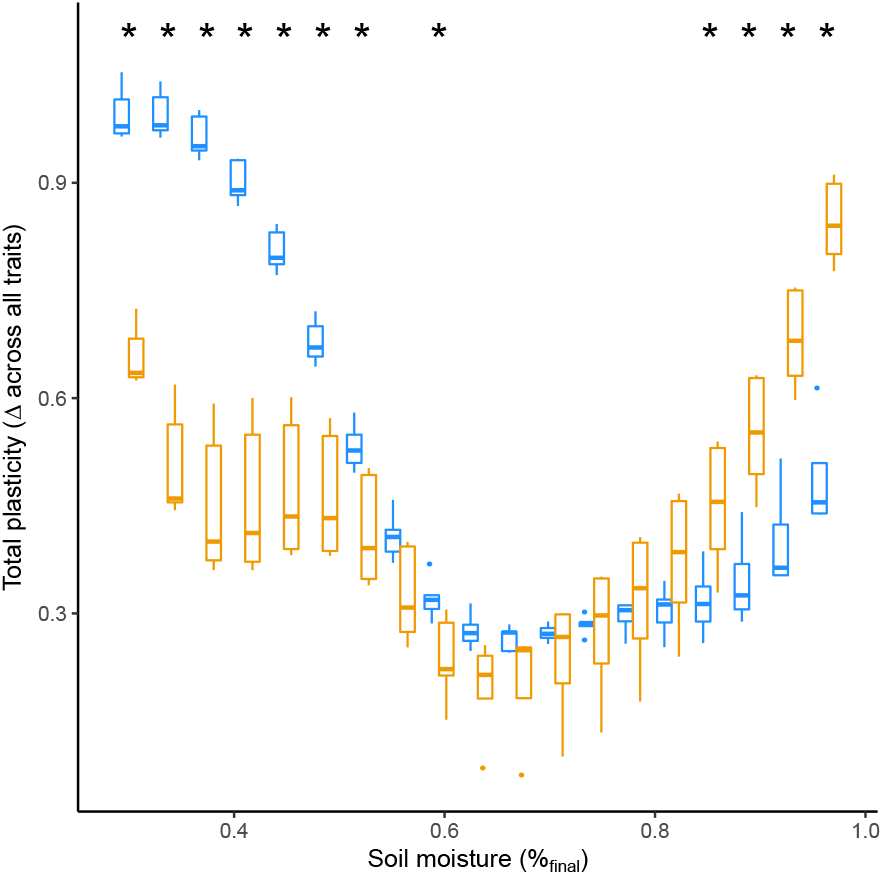
Plasticity through multivariate trait space. (a) Plasticity across all traits was calculated as distance between scaled phenotype for each genotype between different levels of soil moisture. Box plots indicate species median and 25th and 75th percentiles with whiskers extending to 1.5 times the interquartile range. indicates significant differences between species (t-test, *a* < 0.05). *B. sylvaticum* is colored in orange and *B. distachyon* blue.

### Most traits show significant genetic variation

We also tested whether there was significant natural variation for the traits measured between genotypes in each species by looking at the parameters in the final model for each trait. Interestingly, across species, most traits (21/24) included a genotype term in the final model, indicating significant differences between genotypes in magnitude of traits across all levels of soil moisture (Table 1, Figure 2). Though not formally tested, for other traits there are clear distinctions between the two species. For example, *δC_13_* was considerably higher in *B. sylvaticum* (Figure 2). For SLA, while *B. distachyon* showed a strong response to soil moisture, especially under the driest conditions, SLA in *B. sylvaticum* was much less responsive to soil moisture. In contrast, *B. sylvaticum* appears to show a more dramatic response in leaf composition estimated by N content and C:N ratio. We note too that most of the traits included an effect of harvest cohort; this is expected due to difference in age (up to 5 days, 9-11% total) of different harvest cohorts.

### Several traits show interactions between genotype and non-linear responses to soil moisture

Significant interactions between genotype and environmental parameters in a final model indicate the presence of genetic variation for plasticity (GxE) (Via and Lande 1985). For those GxE interactions where the environmental parameter is non-linear, significant GxE indicates genetic variation for the shape of reaction norms. SLA and root mass showed a significant interaction between genotype and soil moisture in *B. distachyon.* In *B. sylvaticum,* RWC, N content, leaf green area, C:N ratio, and root to shoot ratio all showed interactions between genotype and soil moisture (Table 1, Figure 2). In each of these cases, the interaction between genotype and environment involved a non-linear environmental predictor, indicating not only variation for the magnitude of plasticity (i.e. slope) but also variation in the shape of responses.

### Correlations between traits change as a function of soil moisture, often in a non-linear fashion

We calculated correlations between genotype trait means across soil moisture for traits where genotypic differences were observed (i.e. genotype predictor in trait models). Certain traits were tightly correlated regardless of environment. For example, correlations near 1 were observed between biomass and green area in both species across soil moisture. More complex relationships between trait correlations and soil moisture are observed in other trait combinations. For traits with genotype by non-linear environment interactions (Table 1), trait correlations showed non-linear relationships with soil moisture as well. Because more of these interactions were found in *B. sylvaticum* the number of trait combinations showing non-linear relations between correlations and soil moisture appears to be higher than in *B. distachyon* (Figure 4). The correlation between C:N ratio and root:shoot ratio in *B. syl-vaticum*, for example, varied from approximately 0.3 under the wettest environment to approximately −0.7 under the driest environment.

**Figure 4.**
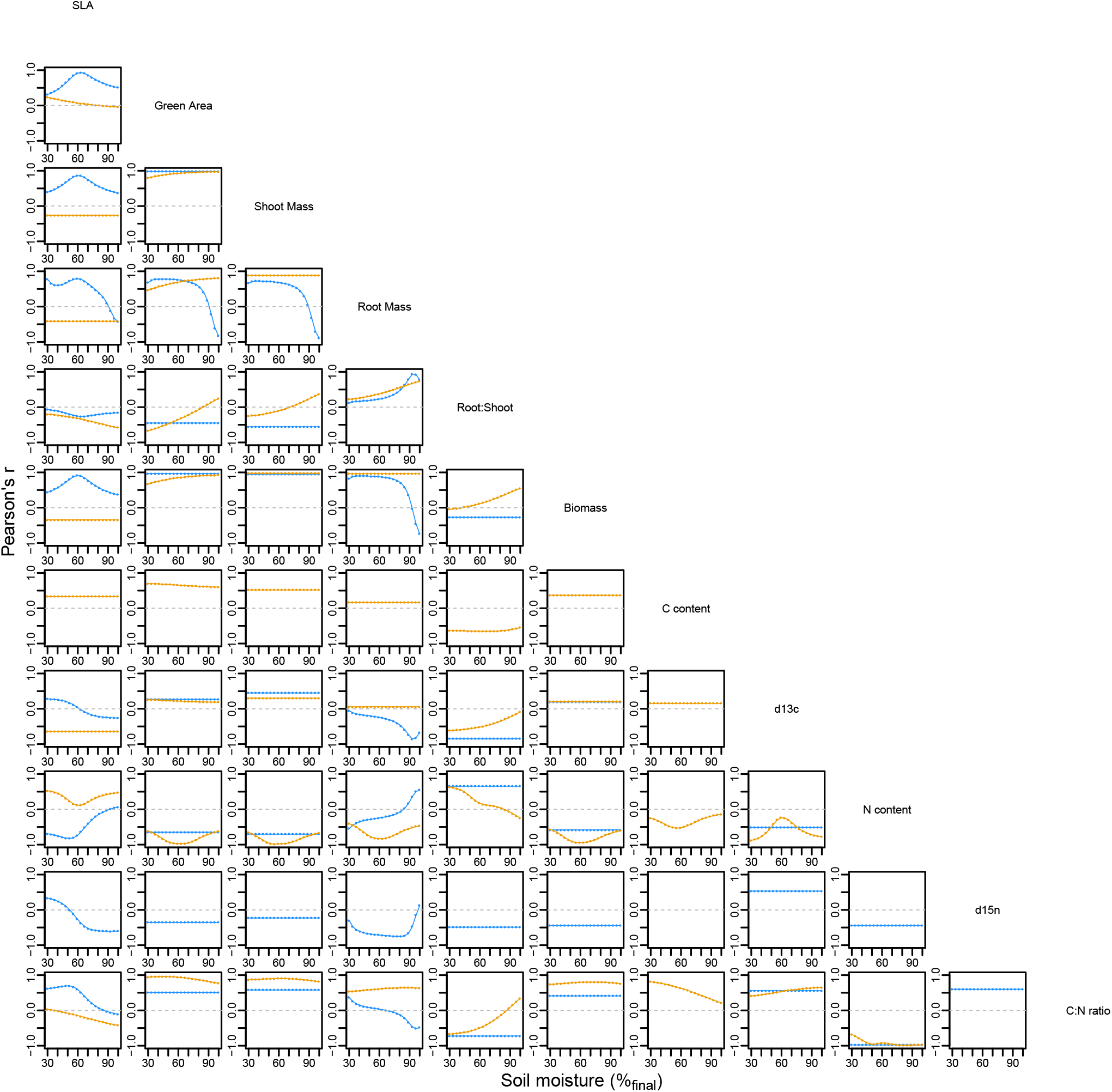
Relationships between trait correlations and soil moisture content. Each point is a correlation coefficient calculated across five genotypes between the intersection of the two traits labeled on the diagonal. Correlations were calculated among genotypes by species (blue = *B. distachyon,* orange = *B. sylvaticum).* Correlations are not shown for traits in species where genotype was not included in final model (Table 1).

**Figure 5.**
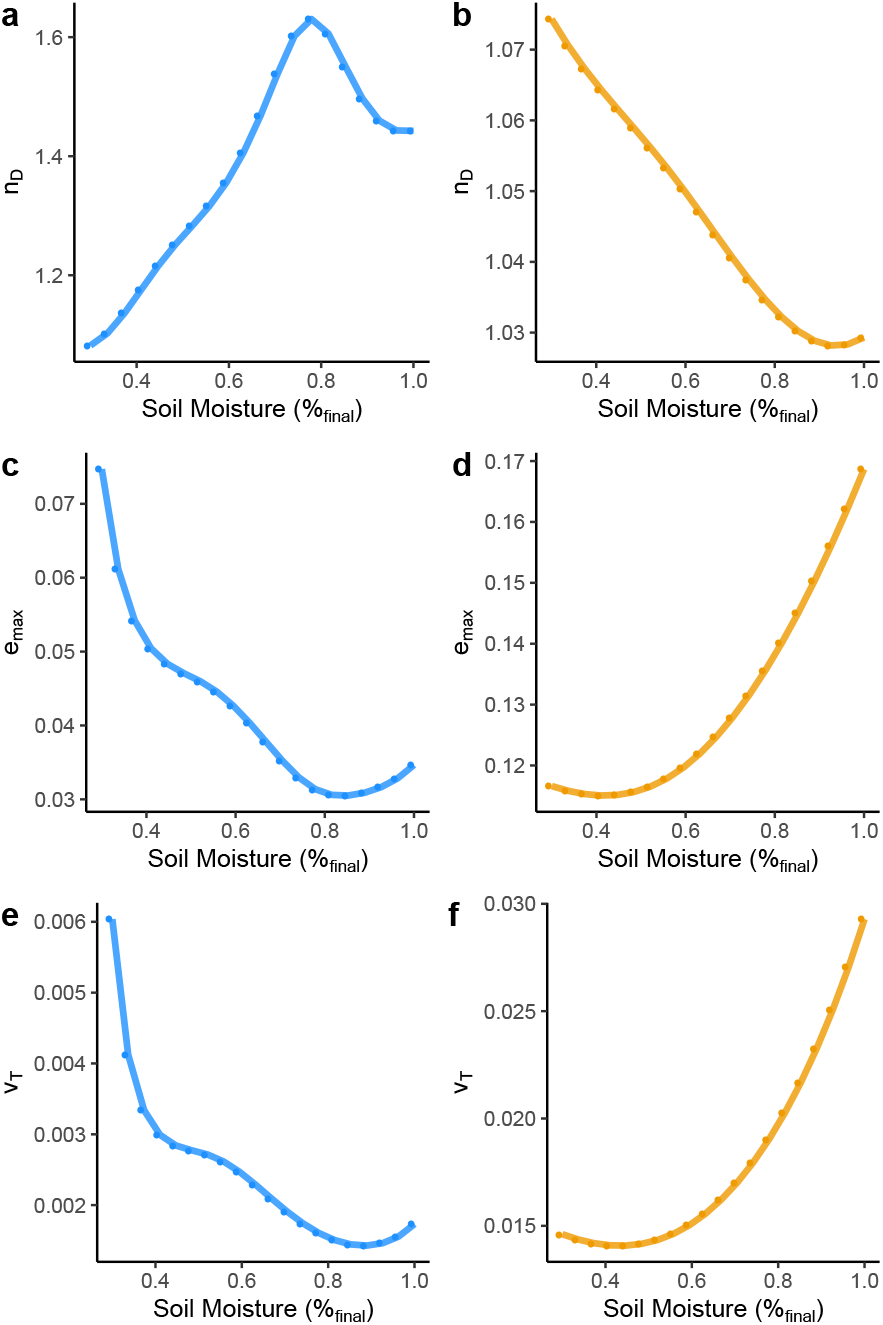
Contrasting patterns of evolutionary constraint between *B. distachyon* and *B. sylvaticum*. Summary statistics of evolutionary constraint as a function of soil moisture in *B. distachyon* (blue) and *B. sylvaticum* (orange). (a and b) The number of effective dimensions, *nD,* estimates number of unconstrained axes of variation. (c and d) The maximum evolvability, *e_max_*, corresponds to the square root of the largest eigenvalue of the genetic covariance matrix. (e and f) The total genetic variance, *v_T_*, is equal to the sum of the eigenvalues of the genetic covariance matrix.

### Evolutionary constraints differ as a function of soil moisture and show contrasting patterns between *Brachypodium* species

To assess evidence of evolutionary constraint on the sampled traits, we approximated and analyzed parameters of the genetic covariance matrix, *G,* in each species across the soil moisture gradient. These analyses revealed contrasting patterns of evolutionary constraint both in relation to soil moisture and between *B. distachyon* and *B. sylvaticum.* In *B. distachyon,* the number of effective dimensions (*n_D_*, which estimates number of axes of variation unconstrained by covariance) was lower when soils were drier. In contrast, *n_D_* was lower in *B. sylvaticum* when soils were wetter. The maximum evolvability *(e_max_,* variance through largest eigenvector of multidimensional trait space) also showed opposite trends between the two species. Whereas in *B. distachyon e_max_* was highest under the driest conditions, in *B. sylvaticum e_max_* was highest under the wettest conditions. The same trend was seen in total genetic variance (*V_T_*, which summarizes all genetic variance through multidimensional trait space).

## Discussion

### Non-linearity in soil moisture response is common in *Brachypodium*

We found significant non-linearity in response to a soil moisture gradient for all measured traits in at least one of the two species sampled. The best-fit function for some traits was quadratic, while other traits showed more complex responses to the environment which were best fit by a spline function. These results offer new insights with respect to the study of plant response to soil drying. By focusing on the curvature of phenotypic response as the explicitly modeled trait, we avoid contrasts of trait values expressed at arbitrary levels of soil water content which may obscure different thresholds of response among the diverse genotypes under study. SLA in *B. distachyon* exhibits this pattern, as two accessions show a threshold-like response in decreasing SLA as soils become drier, and three accessions express their highest SLA at intermediate soil water content. Leaf N content (on a leaf-mass basis) in *B. sylvaticum* likewise shows considerable diversity of response with two accessions expressing their lowest values at intermediate soil water content, one accession expressing its highest leaf N at intermediate soil water content, and one accession showing a nearly linear response along the soil water content gradient.

Non-linearities in trait responses to soil moisture reinforce the need to consider the consequences of extreme weather events when predicting plastic responses, especially when scaling up to investigating the ecosystem effects of individual plant responses to environmental stress (Felton and Smith 2017). As others have previously noted, if organism responses to the environment show non-linear curves then changes in the environment may result in greater or lesser responses than expected based on predictions based on linear response curves alone (Nussey *et al*. 2007; Brommer *et al*. 2008; Visser 2008; Gienapp and Brommer 2014). This suggests, for example, that responses of ecosystem traits such as SLA to increasing aridity and drought may depend on the severity of environ-mental change in a threshold-like fashion. Moreover, because traits like SLA have impacts on broader processes including carbon cycling, carbon models incorporating plastic and evolutionary responses may benefit from information about the (non)-linearity of trait responses to environmental change (Donovan *et al*. 2014; Monroe *et al*. 2018; Walker *et al*. 2019).

### Implications for evolution of *Brachypodium*

Leaf N and SLA are two axes of the classic Leaf Economic Spectrum (Wright *et al*. 2004) and so the contrasting responses of these traits between the annual *B. distachyon* and perennial *B. sylvaticum* may reflect broader differences in their life history strategies. We recently reviewed evidence for physiological, anatomical and developmental differences between herbaceous annual and perennial species, finding support for generally higher SLA in annuals, befitting a generally resource-acquisitive strategy (Lundgren and Des Marais 2020). Garnier (Garnier 1992) argued that small changes in leaf anatomy (e.g. SLA) will likely have large effects on plant growth rate and resource use and could therefore tip the balance between perennial and annual strategies.

We also found that signatures of evolutionary constraint differ along our imposed soil water content gradient. Specifically, we find evidence of highest evolvability in multitrait space (measured by *emax* and *vT*) in *B. distachyon* under the driest soils. In contrast, *B. sylvaticum* exhibited evidence of greater evolvability by the same measures under the highest soil water content studied here. However, *B. distachyon* exhibits evidence of higher constraint caused by genetic covariances (low *nD* under dry conditions) These results indicate that *B. distachyon* has increased genetic variance under dry conditions as compared to wet conditions, but that natural selection may be more constrained to act on this variation due to covariance between traits. In contrast, our results suggest that *B. sylvaticum* has decreased genetic variance under dry conditions on which selection might act but that this variation is less constrained by covariance between traits.

Consistent with predictions of elevated genetic variation revealed in less frequently encountered environments (Gibson and Dworkin 2004; Schlichting 2008; Paaby and Rockman 2014), we speculate that the pattern observed could be a reflection of the different life history strategies of these two species. Annuality is considered a drought adaptive strategy characterized by escaping drought through phenology, by flowering before and remaining dormant as seeds during the most drought prone seasons (Friedman and Rubin 2015; Monroe *et al*. 2019; Friedman 2020). Thus, because of their life history, populations of annuals may actually experience fewer episodes of strong selection from extreme drought, which could explain why we find elevated genetic variance under these environments. In contrast, perennials such as *B. sylvaticum* are subjected to all seasons and might therefore experience more frequent episodes of selection caused by dehydration stress, despite paradoxically being in found in environments where droughts are less frequent on an annual basis. This pattern is consistent with the predictions of cryptic genetic variation revealed under environments where selection is less frequent or severe (Gibson and Dworkin 2004; Schlichting 2008; Paaby and Rockman 2014). We remain cautious about drawing this conclusion from our data, as they represent a comparison between only two species and include a small sample genotypic diversity in each. However, that the difference in patterns of constraint are explained by alternative life history strategies emerges as an interesting hypothesis from these observations, one that might be addressed by future work extending the approaches here to a broader phylogenetic comparison between annual and perennial species.

### Potential to scale up investigations of function-valued responses to soil moisture

We found that genetic correlations between traits can vary over relatively small changes in soil moisture (Fig. 4). Similarly, we found that signatures of evolutionary constraint varied across the environmental gradient, suggesting that responses to selection may be improved or restricted in accordance with patterns of constraint in relation to environment. Interestingly, we also found that some species may be more responsive to selection in a given environment based on patterns of constraint. However, in this study we focus on only five genotypes per species. This reflects the inherent challenge of investigating large numbers of genotypes and high resolution environment gradients. But larger-scale investigations are needed if results are to be extended to the large populations typical of genetic mapping studies and breeding pools. Indeed, responses to selection for drought tolerance may depend on drought severity because of these patterns in genetic correlations. And in the context of breeding, larger studies applying these approaches might be valuable for identifying conditions for which genetic correlations are aligned with breeding objectives (Bernardo 2020).

From a practical perspective this work highlights the potential of function-valued trait approaches that may be scaled up to studying plant-water relations. In this experiment, we investigated variation in plant responses to a gradient of soil moisture using six watering levels, which in combination with random variation in water capacity of experimental pots, produced a continuous gradient of soil moisture ranging from field capacity of the soil to strongly water-limited. In the field, multiple watering regimes in combination with random variability between plots may produce similar gradients of soil moisture. Here, water content was measured gravimetrically. New sensing technologies may be useful for quantifying soil moisture or other continuous environmental gradients in an analogous fashion, to define soil moisture quantitatively and then apply function-valued statistical approaches to contrast trait expression among genotypes. Finally, while in this experiment we used destructive phenotyping methods to measure traits, non-destructive (and high throughput) phenotyping enable measurement of yield or fitness data as well to examine explicit connections between trait variation and adaptation to different degrees of soil moisture in larger experiments. Together these approaches provide opportunities to scale up the analytical framework used here to study genetic variation for soil moisture function-valued-traits in populations of diverse genotypes.

## Acknowledgements

We thank Chase Mason, Eric Goolsby, and Kyle Hernandez for insightful conversations about function-valued trait approaches. We thank Benoit Pujol and two anonymous reviewers for feedback on earlier versions of this manuscript. This work was supported by an Eco-Evo-Devo RCN training grant, USDA-NIFA Award 2014-38420-21801, and Max Planck Society support for JGM. Version 5 of this preprint has been peer-reviewed and recommended by Peer Community In Evolutionary Biology (https://doi.org/10.24072/pci.evolbiol.100119).

## Contributions

JGM and DLD funded, planned, and conducted the experiment. JGM, HC, and DLD contributed to analyses and writing.

## Supplement

**Table S1.**
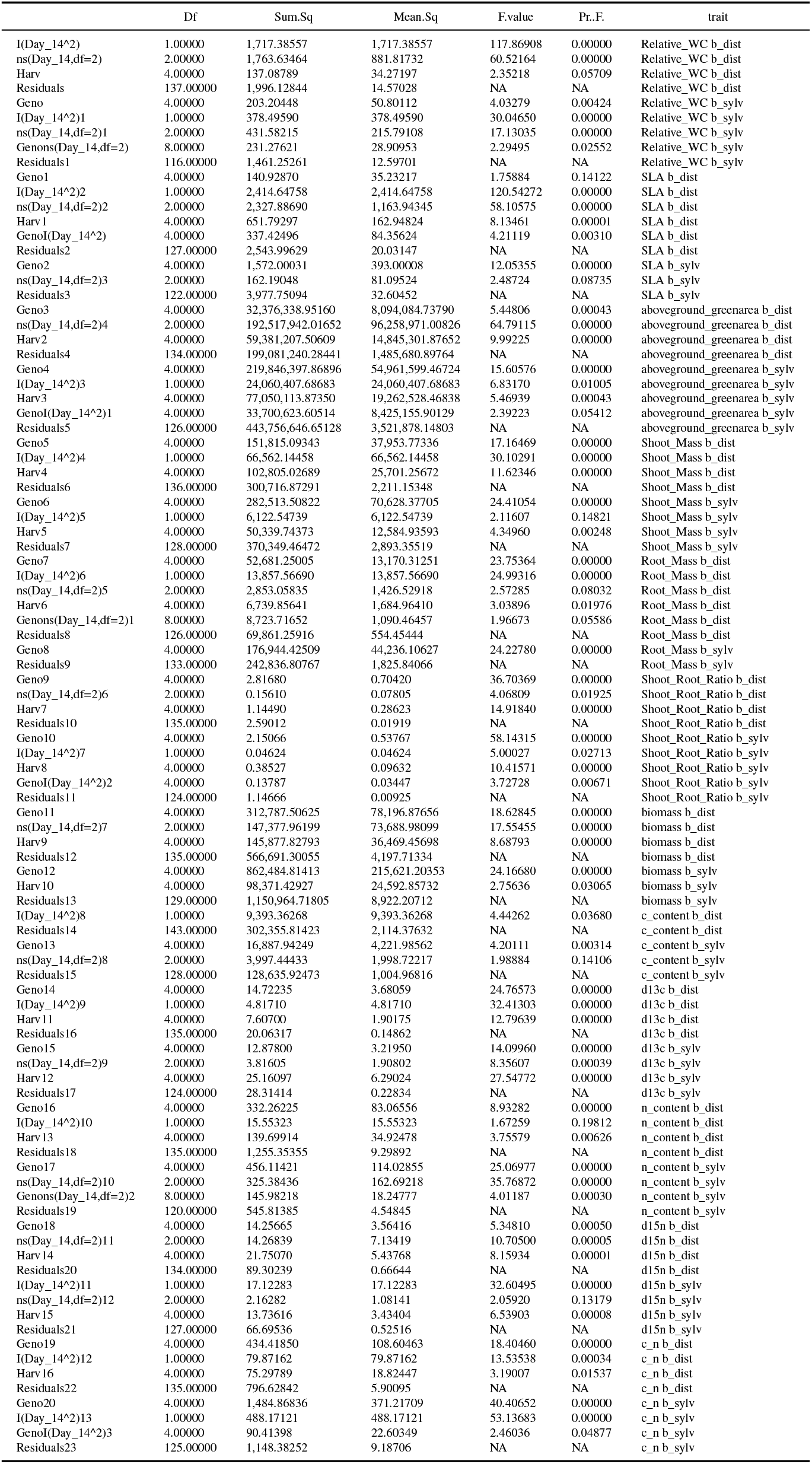
final models

**Table S2.** Selected model r-squared and AIC values

| trait | sp | r2 | AIC |
| --- | --- | --- | --- |
| Relative_WC | b_dist | 0.644 | 809.716 |
| Relative_WC | b_sylv | 0.460 | 725.960 |
| SLA | b_dist | 0.698 | 851.463 |
| SLA | b_sylv | 0.304 | 824.383 |
| aboveground_greenarea | b_dist | 0.588 | 2,484.703 |
| aboveground_greenarea | b_sylv | 0.444 | 2,522.983 |
| Shoot_Mass | b_dist | 0.516 | 1,550.356 |
| Shoot_Mass | b_sylv | 0.478 | 1,503.130 |
| Root_Mass | b_dist | 0.548 | 1,357.246 |
| Root_Mass | b_sylv | 0.422 | 1,434.886 |
| Shoot_Root_Ratio | b_dist | 0.614 | -150.328 |
| Shoot_Root_Ratio | b_sylv | 0.703 | -239.448 |
| biomass | b_dist | 0.517 | 1,644.869 |
| biomass | b_sylv | 0.455 | 1,657.609 |
| c_content | b_dist | 0.030 | 1,525.673 |
| c_content | b_sylv | 0.140 | 1,325.141 |
| d13c | b_dist | 0.575 | 146.704 |
| d13c | b_sylv | 0.596 | 196.255 |
| n_content | b_dist | 0.280 | 746.466 |
| n_content | b_sylv | 0.630 | 603.709 |
| d15n | b_dist | 0.360 | 365.210 |
| d15n | b_sylv | 0.331 | 305.920 |
| c_n | b_dist | 0.425 | 680.522 |
| c_n | b_sylv | 0.642 | 694.126 |

**Figure S1.**
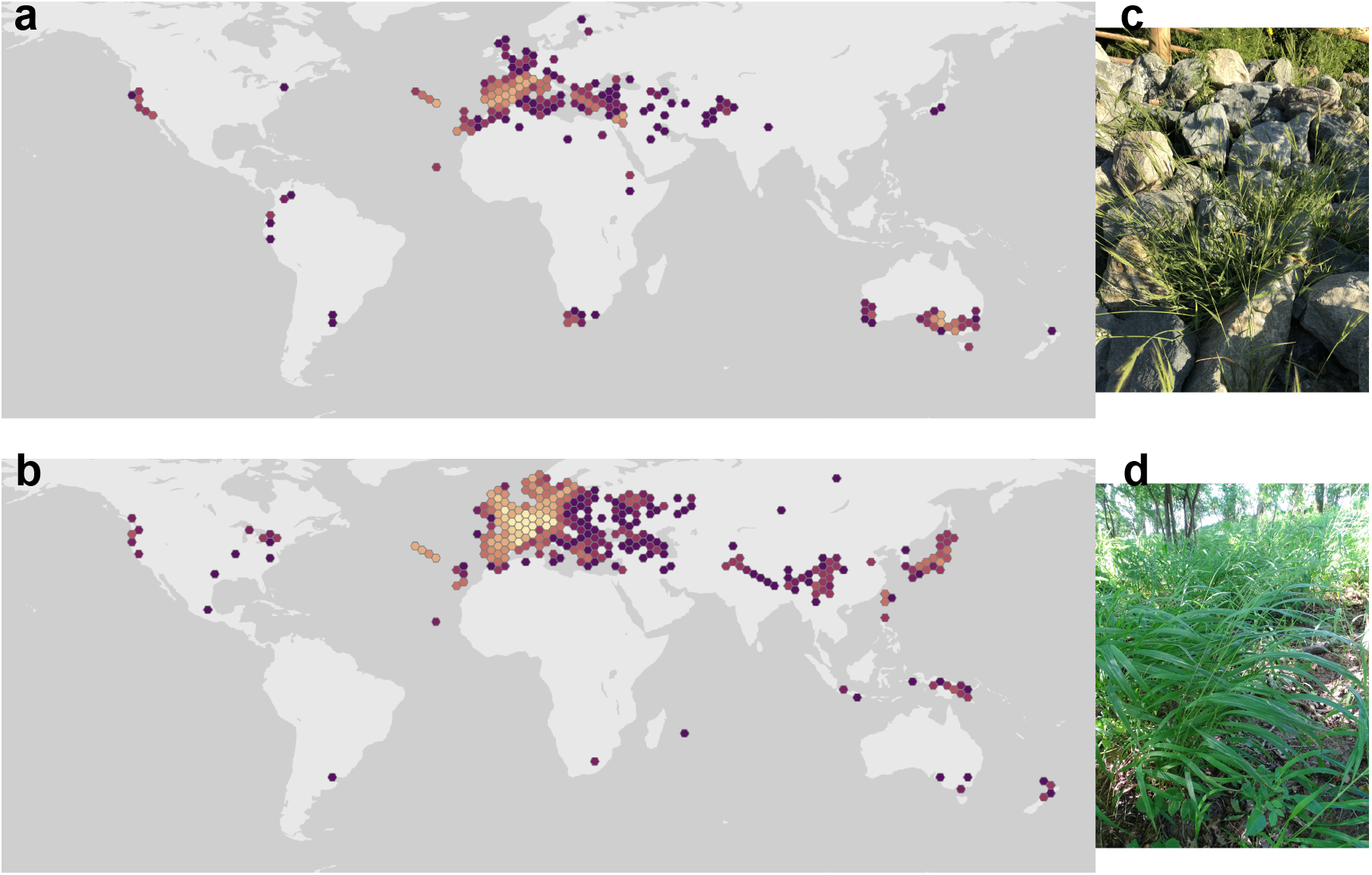
Distributions of (a) *B. distachyon* and (b) *B. sylvaticum* reported on GBIF as of 2019.18.08. Examples of (c) *B. distachyon* Carly Slawson (CC BY 4.0, https://www.inaturalist.org/photos/42532397) and (d) *B. sylvaticum* Grzegorz Grzejszczak (CC BY-NC 4.0, https://www.inaturalist.org/photos/36088991) (GBIF.org (26 February 2020) GBIF Occurrence Download https://doi.org/10.15468/dl.rau5v9).

**Figure S2.**
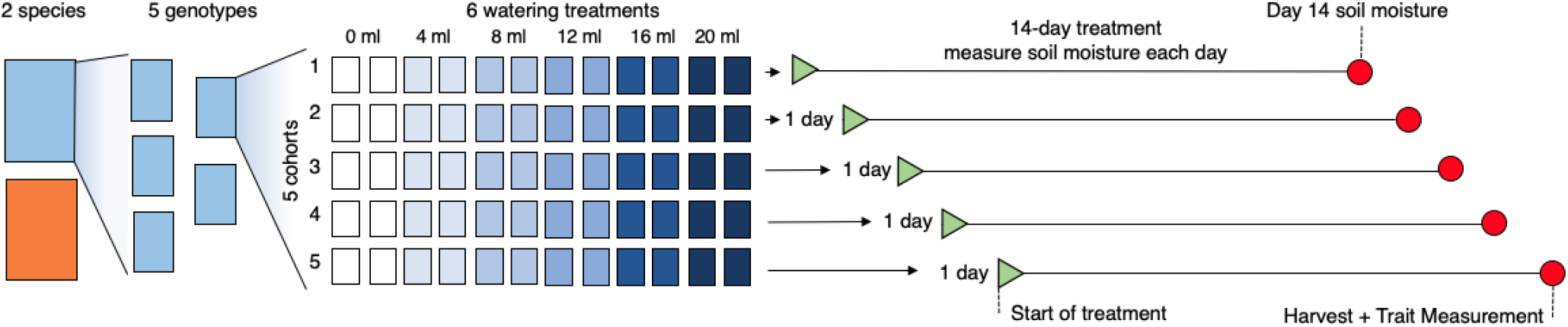
Experimental design. Shown is the schema for each of the 10 genotypes.

**Figure S3.**
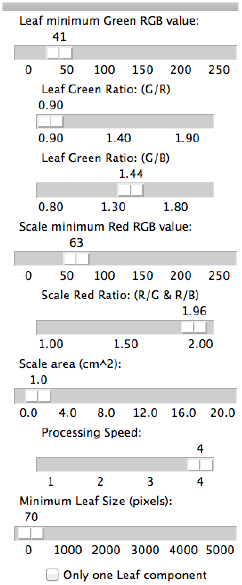
Settings used in Easy Leaf Area.

**Figure S4.**
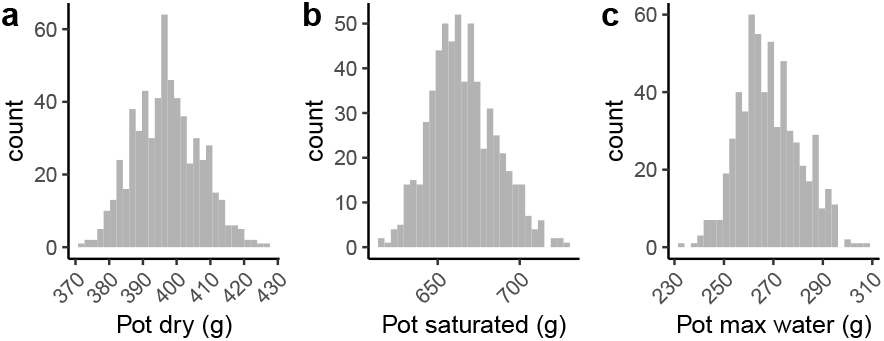
Variation in pot field capacity.

**Figure S5.**
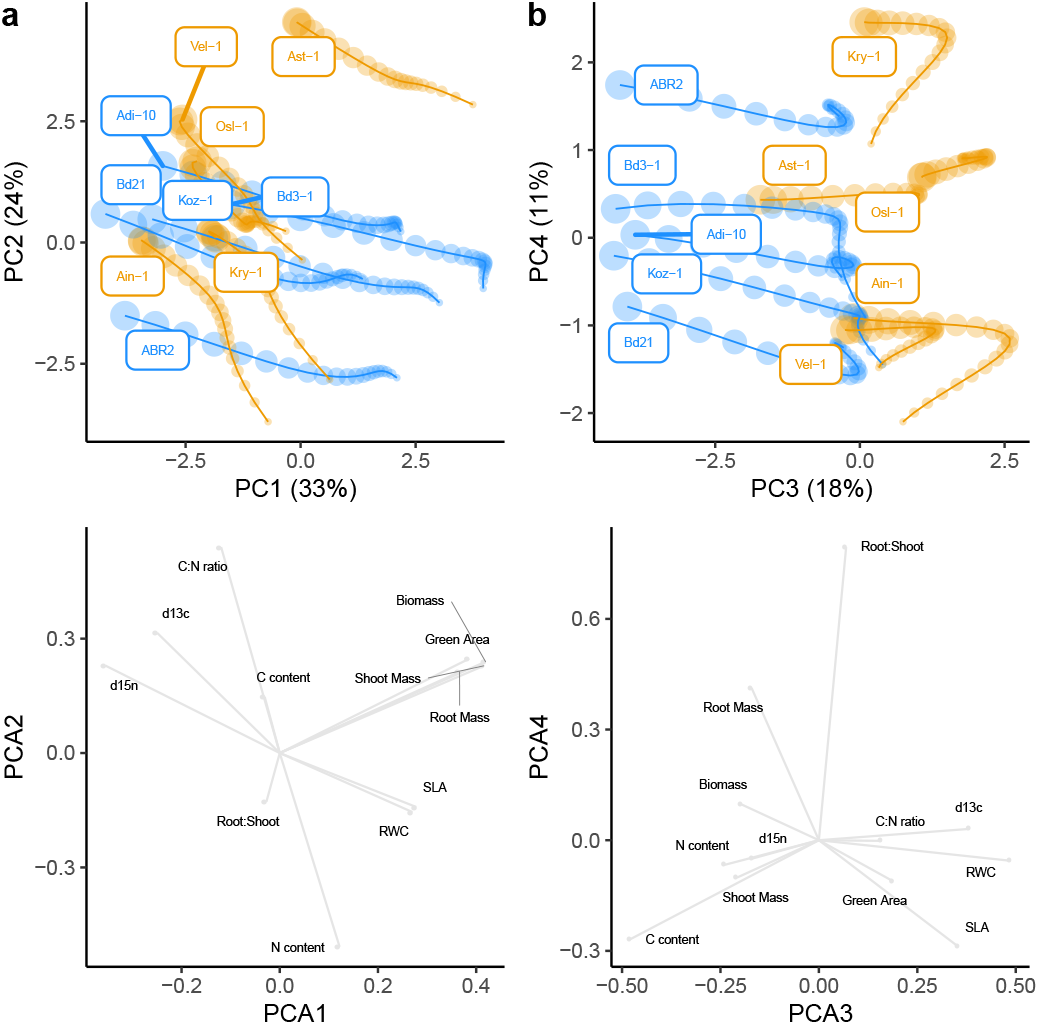
Plasticity through multivariate trait space. Principal component analysis of scaled phenotypic responses to soil moisture gradient among genotypes of both species. Upper panels show genotype means across soil moisture content. Percent values in axis titles indicate percent variance explained by that principal component. Lower panels show eigenvectors of each trait. (a) PC1 and PC2. (b) PC3 and PC4. *B. sylvaticum* is colored in orange and *B. distachyon* blue.

## References

Bernardo R., 2020 Breeding for quantitative traits in plants. Stemma press Woodbury, MN.

Blows M. W., 2007 A tale of two matrices: Multivariate approaches in evolutionary biology. Journal of evolutionary biology 20: 1–8.

Bolstad G. H., W. S. Armbruster, C. Pélabon, R. Pérez-Barrales, and T. F. Hansen, 2010 Direct selection at the blossom level on floral reward by pollinators in a natural population of dalechampia schottii: Fulldisclosure honesty? New Phytologist 370–384.

Brkljacic J., E. Grotewold, R. Scholl, T. Mockler, and D. F. Garvin et al., 2011 Brachypodium as a model for the grasses: Today and the future. Plant Physiology 157: 3–13.

Brommer J. E., K. Rattiste, and A. J. Wilson, 2008 Exploring plasticity in the wild: Laying date-temperature reaction norms in the common gull larus canus. Proceedings of the Royal Society B: Biological Sciences 275: 687–693.

Casper B., I. Forseth, H. Kempenich, S. Seltzer, and K. Xavier, 2001 Drought prolongs leaf life span in the herbaceous desert perennial cryptantha flava. Functional Ecology 15: 740–747.

Catalan P., D. Lopez-Alvarez, A. Diaz-Perez, R. Sancho, and M. L. Lopez-Herranz, 2016 Phylogeny and evolution of the genus brachypodium, in Genetics and genomics of brachypodium, Plant genetics and genomics: Crop models. edited by Vogel J. Springer International.

Des Marais D. L., J. K. McKay, J. H. Richards, S. Sen, and T. Wayne et al., 2012 Physiological genomics of response to soil drying in diverse arabidopsis accessions. The Plant Cell 24: 893–914.

Des Marais D. L., J. R. Lasky, P. E. Verslues, T. Z. Chang, and T. E. Juenger, 2017 Interactive effects of water limitation and elevated temperature on the physiology, development and fitness of diverse accessions of brachypodium distachyon. New Phytologist 214: 132–144.

Dittberner H., A. Korte, T. Mettler-Altmann, A. P. Weber, and G. Monroe et al., 2018 Natural variation in stomata size contributes to the local adaptation of water-use efficiency in arabidopsis thaliana. Molecular ecology 27: 4052–4065.

Donovan L. A., C. M. Mason, A. W. Bowsher, E. W. Goolsby, and C. D. Ishibashi, 2014 Ecological and evolutionary lability of plant traits affecting carbon and nutrient cycling. Journal of Ecology 102: 302–314.

Easlon H. M., and A. J. Bloom, 2014 Easy leaf area: Automated digital image analysis for rapid and accurate measurement of leaf area. Applications in plant sciences 2: 1400033.

Edwards C. E., B. E. Ewers, C. R. McClung, P Lou, and C. Weinig, 2012 Quantitative variation in water-use efficiency across water regimes and its relationship with circadian, vegetative, reproductive, and leaf gas-exchange traits. Molecular Plant 5: 653–668.

El-Soda M., M. P Boer, H. Bagheri, C. J. Hanhart, and M. Koornneef et al., 2014 Genotype-environment interactions affecting preflowering physiological and morphological traits of brassica rapa grown in two watering regimes. Journal of experimental botany 65: 697–708.

Felton A. J., and M. D. Smith, 2017 Integrating plant ecological responses to climate extremes from individual to ecosystem levels. Philosophical Transactions of the Royal Society B: Biological Sciences 372: 20160142.

Friedman J., and M. J. Rubin, 2015 All in good time: Understanding annual and perennial strategies in plants. American journal of botany 102: 497–499.

Friedman J., 2020 The evolution of annual and perennial plant life histories: Ecological correlates and genetic mechanisms. Annual Review of Ecology, Evolution, and Systematics 51: 461–481.

Garnier E., 1992 Growth analysis of congeric annual and perennial grass species. Journal of Ecology 80: 665–675.

Gibert P., B. Moreteau, J. R. David, and S. M. Scheiner, 1998 Describing the evolution of reaction norm shape: Body pigmentation in drosophila. Evolution 52: 1501–1506.

Gibson G., and I. Dworkin, 2004 Uncovering cryptic genetic variation. Nature Reviews Genetics 5: 681–690.

Gienapp P., and J. E. Brommer, 2014 Evolutionary dynamics in response to climate change. Quantitative genetics in the wild 254: 273.

Gomulkiewicz R., J. G. Kingsolver, P. A. Carter, and N. Heckman, 2018 Variation and evolution of function-valued traits. Annual Review of Ecology, Evolution, and Systematics 49: 139–164.

Goolsby E. W., 2015 Phylogenetic comparative methods for evaluating the evolutionary history of function-valued traits. Systematic biology 64: 568–578.

Greenham K., C. R. Guadagno, M. A. Gehan, T. C. Mockler, and C. Weinig et al., 2017 Temporal network analysis identifies early physiological and transcriptomic indicators of mild drought in brassica rapa. Elife 6: e29655.

Griswold C. K., R. Gomulkiewicz, and N. Heckman, 2008 Hypothesis testing in comparative and experimental studies of function-valued traits. Evolution 62: 1229–1242.

Houle D., 1992 Comparing evolvability and variability of quantitative traits. Genetics 130: 195–204.

Juenger T. E., 2013 Natural variation and genetic constraints on drought tolerance. Current opinion in plant biology 16: 274–281.

Kesari R., J. R. Lasky, J. G. Villamor, Des Marais D. L., and Y.-J. C. Chen et al., 2012 Intron-mediated alternative splicing of arabidopsis p5cs1 and its association with natural variation in proline and climate adaptation. Proceedings of the National Academy of Sciences 109: 9197–9202.

Kingsolver J. G., R. Gomulkiewicz, and P. A. Carter, 2001 Variation, selection and evolution of function-valued traits, pp. 87–104 in Microevolution rate, pattern, process, Springer.

Kingsolver J. G., and R. Gomulkiewicz, 2003 Environmental variation and selection on performance curves. Integrative and Comparative Biology 43: 470–477.

Kingsolver J. G., N. Heckman, J. Zhang, P. A. Carter, and J. L. Knies et al., 2015 Genetic variation, simplicity, and evolutionary constraints for function-valued traits. The American Naturalist 185: E166–E181.

Kirkpatrick M., and N. Heckman, 1989 A quantitative genetic model for growth, shape, reaction norms, and other infinite-dimensional characters. Journal of mathematical biology 27: 429–450.

Kirkpatrick M., 2009 Patterns of quantitative genetic variation in multiple dimensions. Genetica 136: 271–284.

Lande R., and S. J. Arnold, 1983 The measurement of selection on correlated characters. Evolution 1210–1226.

Lenk I., L. H. C. Fisher, M. Vickers, A. Akinyemi, and T. Didion et al., 2019 Transcriptional and metabolomic analyses indicate that cell wall properties are associated with drought tolerance in brachypodium distachyon. Int J Mol Sci 20. https://doi.org/10.3390/ijms20071758

Levins R., 1968 Evolution in changing environments: Some theoretical explorations. Princeton University Press.

Lundgren M. R., and Des Marais D. L., 2020 Life history variation as a model for understanding trade-offs in plant-environment interactions. Current Biology 30: R180–R189.

Luo N., X. Yu, G. Nie, J. Liu, and Y. Jiang, 2016 Specific peroxidases differentiate brachypodium distachyon accessions and are associated with drought tolerance traits. Ann Bot 118: 259–70. https://doi.org/10.1093/aob/mcw104

Mason C. M., M. C. LaScaleia, De La Pascua D. R., J. G. Monroe, and E. W. Goolsby, 2020 Learning from dynamic traits: Seasonal shifts yield insights into ecophysiological trade-offs across scales from macroevolutionary to intraindividual. International Journal of Plant Sciences 181: 88–102.

McGuigan K., 2009 Condition dependence varies with mating success in male drosophila bunnanda. Journal of evolutionary biology 22: 1813–1825.

McGuigan K., N. Nishimura, M. Currey, D. Hurwit, and W. A. Cresko, 2010 Quantitative genetic variation in static allometry in the threespine stickleback. Integrative and comparative biology 50: 1067–1080.

Meyer K., 2005 Random regression analyses using b-splines to model growth of australian angus cattle. Genet. Sel. Evol. 37: 473–500.

Monroe J. G., D. W. Markman, W. S. Beck, A. J. Felton, and M. L. Vahsen et al., 2018 Ecoevolutionary dynamics of carbon cycling in the anthropocene. Trends in ecology & evolution 33: 213–225.

Monroe J. G., B. Gill, K. G. Turner, and J. K. McKay, 2019 Drought regimens predict life history strategies in heliophila. New Phytologist 223: 2054–2062.

Nussey D., A. Wilson, and J. Brommer, 2007 The evolutionary ecology of individual phenotypic plasticity in wild populations. Journal of evolutionary biology 20: 831–844.

Paaby A. B., and M. V. Rockman, 2014 Cryptic genetic variation: Evolution’s hidden substrate. Nature Reviews Genetics 15: 247–258.

Passioura J., 1996 Drought and drought tolerance. Plant growth regulation 20: 79–83.

Pearse I. S., J. M. Aguilar, and S. Y. Strauss, 2019 Life history plasticity and water use trade-offs associated with drought resistance in a clade of california jewelflowers

Pettay J. E., A. Charmantier, A. J. Wilson, and V. Lummaa, 2008 Age-specific genetic and maternal effects in fecundity of preindustrial finnish women. Evolution: International Journal of Organic Evolution 62: 2297–2304.

Robinson M. R., A. J. Wilson, J. G. Pilkington, T. H. Clutton-Brock, and J. M. Pemberton et al., 2009 The impact of environmental heterogeneity on genetic architecture in a wild population of soay sheep. Genetics 181: 1639–1648.

Rocha F. B., and L. B. Klaczko, 2012 Connecting the dots of nonlinear reaction norms unravels the threads of genotype-environment interaction in drosophila. Evolution: International Journal of Organic Evolution 66: 3404–3416.

Schlichting C. D., 2008 Hidden reaction norms, cryptic genetic variation, and evolvability. Annals of the New York Academy of Sciences 1133: 187–203.

Skirycz A., H. Claeys, De Bodt S., A. Oikawa, and S. Shinoda et al., 2011 Pause-and-stop: The effects of osmotic stress on cell proliferation during early leaf development in arabidopsis and a role for ethylene signaling in cell cycle arrest. Plant Cell 23: 1876–88. https://doi.org/10.1105/tpc.111.084160

Steinwand M. A., H. A. Young, J. N. Bragg, C. M. Tobias, and J. P Vogel, 2013 Brachypodium sylvaticum, a model for perennial grasses: Transformation and inbred line development. PLoS One 8: e75180. https://doi.org/10.1371/journal.pone.0075180

Stinchcombe J. R., R. Izem, M. S. Heschel, B. V. McGoey, and J. Schmitt, 2010 Across-environment genetic correlations and the frequency of selective environments shape the evolutionary dynamics of growth rate in impatiens capensis. Evolution: International Journal of Organic Evolution 64: 2887–2903.

Stinchcombe J. R., M. Kirkpatrick, F.-v. T. W. Group, and others, 2012 Genetics and evolution of function-valued traits: Understanding environmentally responsive phenotypes. Trends in Ecology & Evolution 27: 637–647.

Van Eeuwijk F., 1995 Linear and bilinear models for the analysis of multi-environment trials: I. An inventory of models. Euphytica 84: 1–7.

Vasseur F., T. Bontpart, M. Dauzat, C. Granier, and D. Vile, 2014 Multivariate genetic analysis of plant responses to water deficit and high temperature revealed contrasting adaptive strategies. Journal of experimental botany 65: 6457–6469.

Venables W. N., and B. D. Ripley, 2002 Modern applied statistics with s. Springer, New York.

Verelst W., E. Bertolini, De Bodt S., K. Vandepoele, and M. Demeulenaere et al., 2013 Molecular and physiological analysis of growth-limiting drought stress in brachypodium distachyon leaves. Mol Plant 6: 311–22. https://doi.org/10.1093/mp/sss098

Verslues P E., and T. E. Juenger, 2011 Drought, metabolites, and arabidopsis natural variation: A promising combination for understanding adaptation to water-limited environments. Current opinion in plant biology 14: 240–245.

Via S., and R. Lande, 1985 Genotype-environment interaction and the evolution of phenotypic plasticity. Evolution 39: 505–522.

Visser M. E., 2008 Keeping up with a warming world; assessing the rate of adaptation to climate change. Proceedings of the Royal Society B: Biological Sciences 275: 649–659.

Vogel J. P., M. Tuna, H. Budak, N. Huo, and Y. Q. Gu et al., 2009 Development of ssr markers and analysis of diversity in turkish populations of brachypodium distachyon. BMC Plant Biology 9: 88.

Walker T. W., W. Weckwerth, L. Bragazza, L. Fragner, and B. G. Forde et al., 2019 Plastic and genetic responses of a common sedge to warming have contrasting effects on carbon cycle processes. Ecology letters 22: 159–169.

Wood C. W., and Brodie III E. D., 2015 Environmental effects on the structure of the g-matrix. Evolution 69: 2927–2940.

Wright I. J., P B. Reich, M. Westoby, D. D. Ackerly, and Z. Baruch et al., 2004 The worldwide leaf economics spectrum. Nature 428: 821.

Yarkhunova Y., C. E. Edwards, B. E. Ewers, R. L. Baker, and T. L. Aston et al., 2016 Selection during crop diversification involves correlated evolution of the circadian clock and ecophysiological traits in brassica rapa. New Phytologist 210: 133–144.

Bernardo R., 2020 Breeding for quantitative traits in plants. Stemma press Woodbury, MN.

Passioura J., 1996 Drought and drought tolerance. Plant growth regulation 20: 79–83.

Venables W. N., and B. D. Ripley, 2002 Modern applied statistics with s. Springer, New York.

Wood C. W., and Brodie IIIE. D., 2015 Environmental effects on the structure of the g-matrix. Evolution 69: 2927–2940.

Wright I. J., P. B. Reich, M. Westoby, D. D. Ackerly, and Z. Baruch et al., 2004 The worldwide leaf economics spectrum. Nature 428: 821.

